# Recent and rapid colonization of the Lesser Sundas Archipelago from adjacent Sundaland by seven amphibian and reptile species

**DOI:** 10.1101/571471

**Authors:** Sean B. Reilly, Alexander L. Stubbs, Benjamin R. Karin, Evy Arida, Djoko T. Iskandar, Jimmy A. Mcguire

## Abstract

The Lesser Sundas Archipelago is comprised of two parallel chains of islands that extend between the Asian continental shelf (Sundaland) and Australo-Papuan continental shelf (Sahul). These islands have served as stepping-stones for taxa dispersing between the Asian and Australo-Papuan biogeographic realms. While the oceanic barriers have prevented many species from colonizing the archipelago, a number of terrestrial vertebrate species have colonized the islands either by rafting/swimming or human introduction. Here we examine phylogeographic structure within the Lesser Sundas for three snake, two lizard, and two frog species that each have a Sunda Shelf origin. These species are suspected to have recently colonized the archipelago, though all have inhabited the Lesser Sundas for over 100 years. We sequenced mtDNA from 230 samples to test whether there is sufficiently deep genetic structure within any of these taxa to reject human-mediated introduction. Additionally, we tested for genetic signatures of population expansion consistent with recent introduction, and estimated the ages of Lesser Sundas clades, if any exist. Our results show little to no genetic structure between populations on different islands in five species, and moderate structure in two species. Nucleotide diversity is low for all species, and the ages of the most recent common ancestor for species with monophyletic Lesser Sundas lineages date to the Holocene or late Pleistocene. These results support the hypothesis that these species entered the archipelago relatively recently and either naturally colonized or were introduced by humans to most of the islands within the archipelago within a short time span.

## 1 INTRODUCTION

The oceanic islands of Wallacea are united by their historical isolation from the land masses of the Sunda Shelf to the west and the Sahul Shelf to the east and south (Figure 1). While oceanic islands tend to have lower biodiversity than adjacent continental regions, they also tend to have a higher proportion of endemic species in part because terrestrial fauna must cross oceanic barriers in order to colonize them making successful colonization rare and gene flow to and from the island low (Whittaker & Fernández-Palacios, 2007). Many other factors also influence the species diversity on oceanic islands such as their distance from continental sources, the presence or absence of intervening stepping-stone islands, island size, habitat heterogeneity, elevation, latitude, age, etc. (MacArthur & Wilson, 2001; Whittaker & Fernández-Palacios, 2007). However, since humans began maritime travel, the accumulation of species by human-mediated introduction has had an immense impact on the species diversity of islands worldwide (see Austin, 1999; Caphina *et al.*, 2015).

**FIGURE 1.**
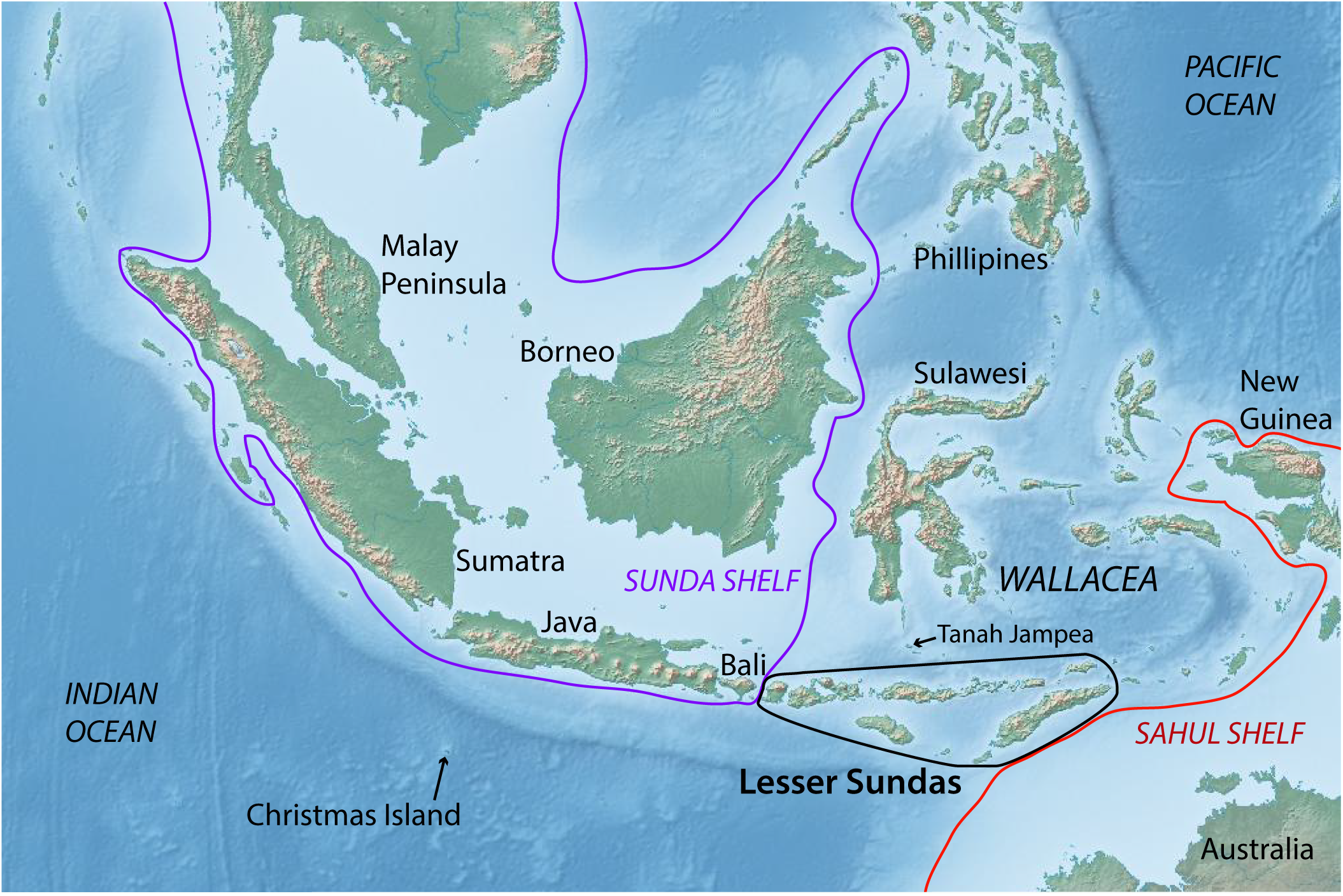
Map of the Sunda Shelf and Wallacea regions with the borders of the Sunda Shelf shown as a yellow line and the border of the Sahul Shelf shown as a red line. Wallacea consists of the oceanic islands between those two shelves.

The Lesser Sundas Archipelago, comprising the southern portion of Wallacea, is composed of two parallel, linearly-arranged chains of oceanic islands, the oldest of which have been continuously emergent for 10-12 Ma (Hall, 2009). This long period of isolation provided ample time for natural colonization of the islands, as well as for *in situ* diversification, and the Lesser Sundas are consequently home to many endemic species (Orne *et al*., 2005). Recent studies of Lesser Sundas amphibians and reptiles have shown that many of these endemics can be expected to exhibit deep inter-and intra-island divergences that reflect the complex tectonic history of the archipelago (Reilly, 2016; Reilly *et al.*, 2019). Although much of the diversity of the Lesser Sundas originated via natural processes, the archipelago has been inhabited by sea faring humans for over 40,000 years, and these humans maintained a long tradition of pelagic fishing and trade between islands (O’Connor *et al.*, 2011). Indonesia is currently the fourth most populous country on earth and traveling and movement of goods between islands is still commonly undertaken by boat (Monk *et al.*, 1997). The long period of habitation by sea-faring humans has resulted in substantial human-mediated dispersal of plants and animals, including many reptile and amphibian species, throughout the archipelago (Heinsohn, 2003; Reilly *et al.*, 2017).

Many species of reptiles and amphibians have become highly invasive due to human-mediated dispersal with major impacts on native fauna (Kraus, 2015). In the Lesser Sundas, this phenomenon is easily observable by simply boarding one of the many ferries that traverse the archipelago, upon which multiple species of small geckos can be seen crawling on the walls of the ship. However, there are a number of other species of reptiles and amphibians in the Lesser Sundas with colonization histories that remain unclear. Here we examine seven species of reptiles and amphibians, each occurring on most of the major islands (Figure 2), that have either been shown to be recently introduced into other regions or are suspected of being moved between islands in the Lesser Sundas (Heinsohn, 2003). Nevertheless, these seven species have occupied islands within the archipelago for at least 100 years, indicating that they either arrived via human-mediated introduction before the early 1900’s (see van Lidth de Jeude, 1895; Boulenger, 1897; Barbour, 1912; De Rooij, 1917a, 1917b; van Kampen, 1923; Mertens, 1930), or that they arrived via natural colonization.

**FIGURE 2.**
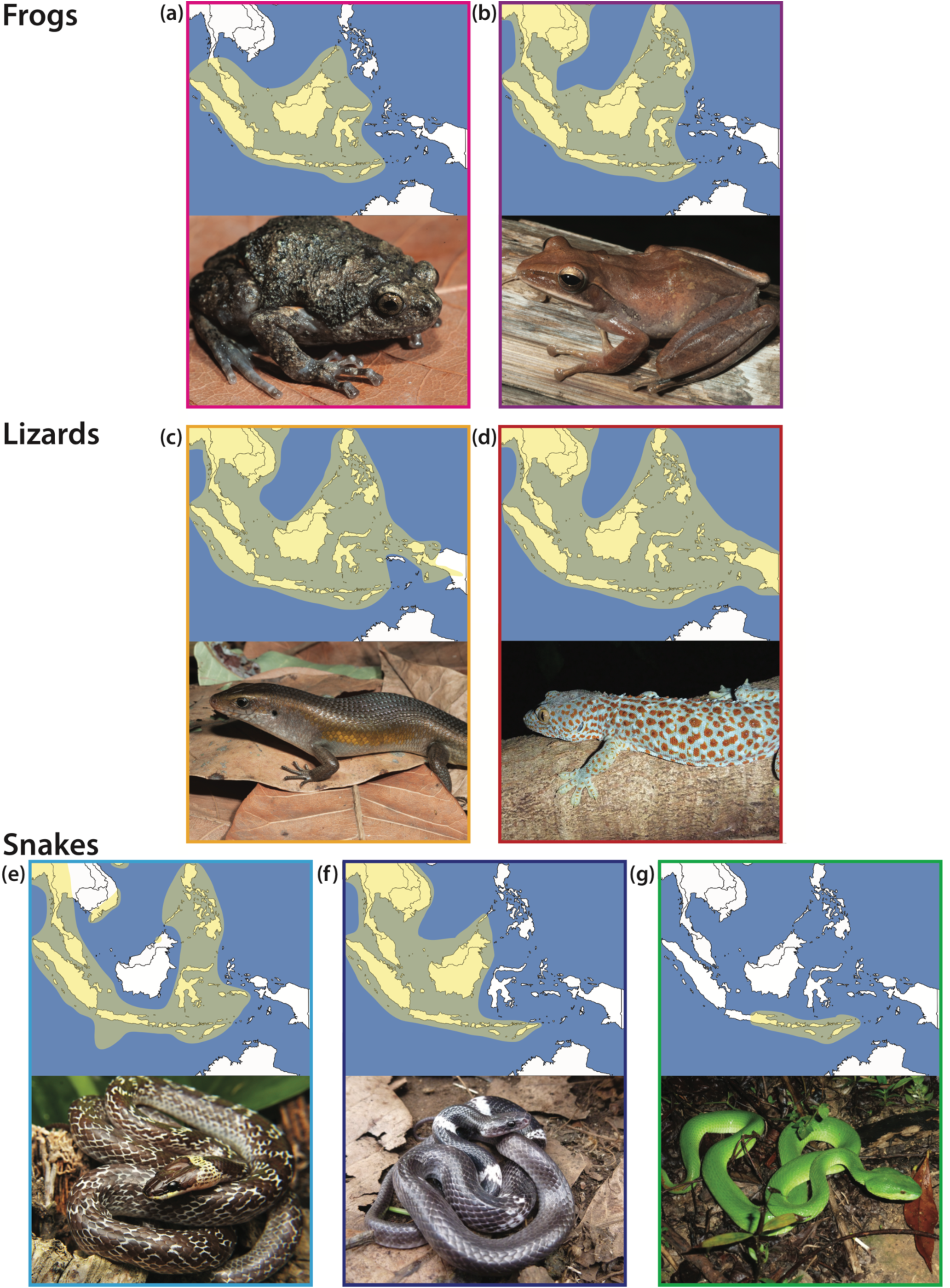
A-G. Photos of focal taxa and their geographic range within Southeast Asia. -A. *Kaloula baleata*, -B. *Polypedates leucomystax*, -C. *Eutropis multifasciata*, -D. *Gekko gecko*, -E. *Lycodon capucinus*, -F. *Lycodon subcinctus*, and -G. *Trimeresurus insularis.*

Here we utilize mtDNA sequence data from 230 newly sequenced samples, along with existing data, to examine the phylogeographical structure in these seven species of reptiles and amphibians with the goal of determining if there is sufficiently deep genetic structure within the Lesser Sundas to reject the hypothesis that they were introduced by humans. We pursue this goal by estimating summary statistics useful for the detection of recent population expansions consistent with recent introductions, estimating the age of monophyletic Lesser Sundas lineages, and generating phylogenies that can shed light on the biogeographical history and any possible phylogeographic structure within the archipelago.

## 2 MATERIALS & METHODS

### 2.1 Focal taxa

*Lycodon capucinus* (Family: Colubridae). The common wolf snake has been shown to be invasive to many regions of Southeast Asia (Fritts, 1993; O’Shea *et al.*, 2018). Because sampling of this species within Indonesia is sparse, it remains unclear whether *L. capucinus* occurs naturally in the Lesser Sunda Islands.

*Lycodon subcinctus* (Family: Colubridae). The white-banded wolf snake also occurs throughout the Lesser Sundas and is similar in size and ecology to *L. capucinus*, suggesting it may have been transported between islands by human activity.

*Trimeresurus insularis* (Family: Viperidae). The venomous white-lipped island pitviper is widespread throughout the archipelago and exhibits regional color morphs that are consistent with long term isolation (see de Lang, 2011; personal observation). However, limited genetic analyses have found low divergence among island populations (How *et al.*, 1996; David *et al.*, 2003; Malhotra & Thorpe, 2004) suggesting the possibility of recent colonization or recent movement between islands.

*Gekko gecko* (Family: Gekkonidae). Tokay geckos occur throughout the Indo-Australian archipelago and a study based on limited sampling found minimal genetic divergence between Timor and multiple Sunda Shelf localities (Roesler *et al.*, 2011). This species is commonly found in human settlements and its prey, insects and smaller geckos, are abundant on boats travelling between islands.

*Eutropis multifasciata* (Family: Scincidae). Sun skinks were shown to be recently introduced to other regions of eastern Indonesia, and even to Australia and the USA (Ingram, 1987; Meshaka *et al.*, 2004; O’Shea *et al.*, 2018). A study based on allozyme data found limited divergence between islands within the Lesser Sundas suggesting that inter-island movement may be prevalent (Schmitt *et al.*, 2000). *Eutropis multifasciata* from the nearest Sunda Shelf island, Bali, was once considered a distinct subspecies (*E. m. balinensis*) from Lesser Sundas populations (*E. m. multifasciata*) based on scale morphology and coloration, though these subspecies have since been synonymized (Mertens, 1927, 1930; Auffenberg, 1980).

*Polypedates leucomystax* (Family: Rhacophoridae). The common tree frog has been shown to be introduced to various parts of Indonesia and the Philippines (Brown *et al*., 2010). However, relatively high genetic diversity on neighboring Java combined with limited sampling from the Lesser Sundas has left their colonization history in the Lesser Sunda Islands an open question (Kuraishi *et al.*, 2013).

*Kaloula baleata* (Family: Microhylidae). The smooth-fingered narrow-mouthed frog is common throughout the Sunda Shelf and the Lesser Sundas. The close relative *K. pulchra* is suspected to have been introduced to multiple islands in Wallacea including Flores, suggesting that *K. baleata* may also have been introduced via human activity (Whitten *et al.*, 1987; Heinsohn, 2003).

### 2.2 Sampling

Herpetological surveys were conducted over the course of five expeditions between 2010-2014 on the islands of Bali, Nusa Penida, Lombok, Sumbawa, Flores, Lembata, Pantar, Alor, Wetar, Timor, Rote, Savu, and Sumba. Sampling information for each species can be found in Table 1 and a list of all newly sequenced samples and their localities can be found in Table S1. Voucher specimens and tissue samples are housed at either the UC Berkeley Museum of Vertebrate Zoology (MVZ) or the Museum Zoologicum Bogoriense (MZB) in Cibinong, Indonesia.

**TABLE 1.** Mitochondrial marker information. bp= base pairs; n=number of samples; LS=includes only samples from Lesser Sundas.

| Species | gene | bp | seq.<br>evol.<br>model | n<br>(LS) | # LS<br>Islands<br>Sampled | LS<br>Variable<br>Sites | LS<br>Parsimony<br>Inf. Sites |
| --- | --- | --- | --- | --- | --- | --- | --- |
| *Kaloula baleata* | *16S* | 831 | GTR | 13 | 4 | 22 | 10 |
| *Polypedates leucomystax* | *16S* | 892 | HKY+I | 41 | 5 | 4 | 0 |
| *Eutropis multifasciata* | *ND4* | 892 | HKY+I | 36 | 8 | 54 | 10 |
| *Gekko gecko* | *ND4* | 840 | HKY | 34 | 10 | 17 | 11 |
| *Lycodon capucinus* | *cytb* | 1,073 | GTR+I | 19 | 8 | 2 | 1 |
| *Lycodon subcinctus* | *cytb* | 1,078 | GTR+I | 8 | 5 | 1 | 0 |
| *Trimeresurus insularis* | *ND4+cytb* | 1,974 | HKY+I | 57 | 9 | 10 | 2 |

### 2.3 Data collection

Genomic DNA was extracted from liver or tail tip tissues using the Qiagen DNeasy blood and tissue kit (Qiagen, Valencia, CA, USA). For *T. insularis*, two mitochondrial genes, *ND4* and *cytochrome b*, were PCR-amplified following standard procedures using the primers ND4 and LEU for the *ND4* gene (Arévalo *et al.* 1994), and the primers THRSN2 and H14910 for the cytochrome b (*cytb*) gene (Burbrink *et al.* 2000). The *cytb* gene was sequenced for both species of *Lycodon* using the primers THRSN2 and H14910. The *ND4* gene was sequenced for both *E. multifasciata* and *G. gecko* using the primers ND4 and LEU. The *16S* gene was sequenced for *P. leucomystax* and *K. baleata* using the primers 16sc-L and 16sd-H (Evans *et al.*, 2003). PCR reactions were carried out using standard Sanger sequencing methods, and ethanol precipitation. DNA sequence visualization was performed on an ABI 3730 automated sequencer (Applied Biosystems, Foster City, CA, USA). Forward and reverse sequence reads were combined in GENEIOUS V11.1.5 (https://www.geneious.com). For each species, relevant DNA sequence data (i.e., for samples from the Lesser Sundas or other Sunda Shelf localities) were downloaded from GenBank. Sequence alignments were generated using MUSCLE (Edgar, 2004) with manual corrections where necessary.

### 2.4 Data analysis

The software JMODELTEST 2 (Darriba *et al.*, 2012) was used to infer the appropriate models of sequence evolution for phylogenetic analyses. Maximum Likelihood phylogenetic analysis was carried out using the program RAXML v8 (Stamatakis, 2014) with node support assessed with 1,000 nonparametric bootstrap replicates. For monophyletic Lesser Sundas clades with phylogenetic structure, the timing of entry into the archipelago was roughly estimated using the software BEAST v2.4.8 (Bouckaert *et al.*, 2014) with a strict molecular clock applied and a rule of thumb 2% divergence/million years sequence evolution rate-calibration. For BEAST analyses, a data matrix composed of one sequence per unique haplotype was subjected to two separate runs of 10 million generations or more (in order to ensure parameter ESS values were >200). To obtain divergence estimates, the results of each pair of analyses were combined after discarding 10% of the samples as burn-in.

Summary statistics including number of haplotypes, parsimony informative sites, haplotype diversity, nucleotide divergence, Tajima’s *D* (Tajima, 1989), and Fu’s *Fs* were calculated with the software DNASP V5 (Librado & Rozas, 2009). Tajima’s *D* (when significantly negative) and Fu’s *Fs* statistics can detect a rapid population expansion after a genetic bottleneck. All sequences are deposited in GenBank (accession numbers available upon acceptance).

## 3 RESULTS

### 3.1 Genetic structure

*Kaloula baleata* exhibits substantial genetic structure (Fig. 3a). Samples from Timor are nested within samples from Java and Bali. This “Java/Bali/Timor” clade is sister to a Lesser Sundas clade containing samples from Sumbawa, Sumba, and Flores. The haplotype of the Bali sample is identical to that of two samples from Timor. Frogs from Sumbawa form a monophyletic assemblage with high support (bootstrap=94).

**FIGURE 3.**
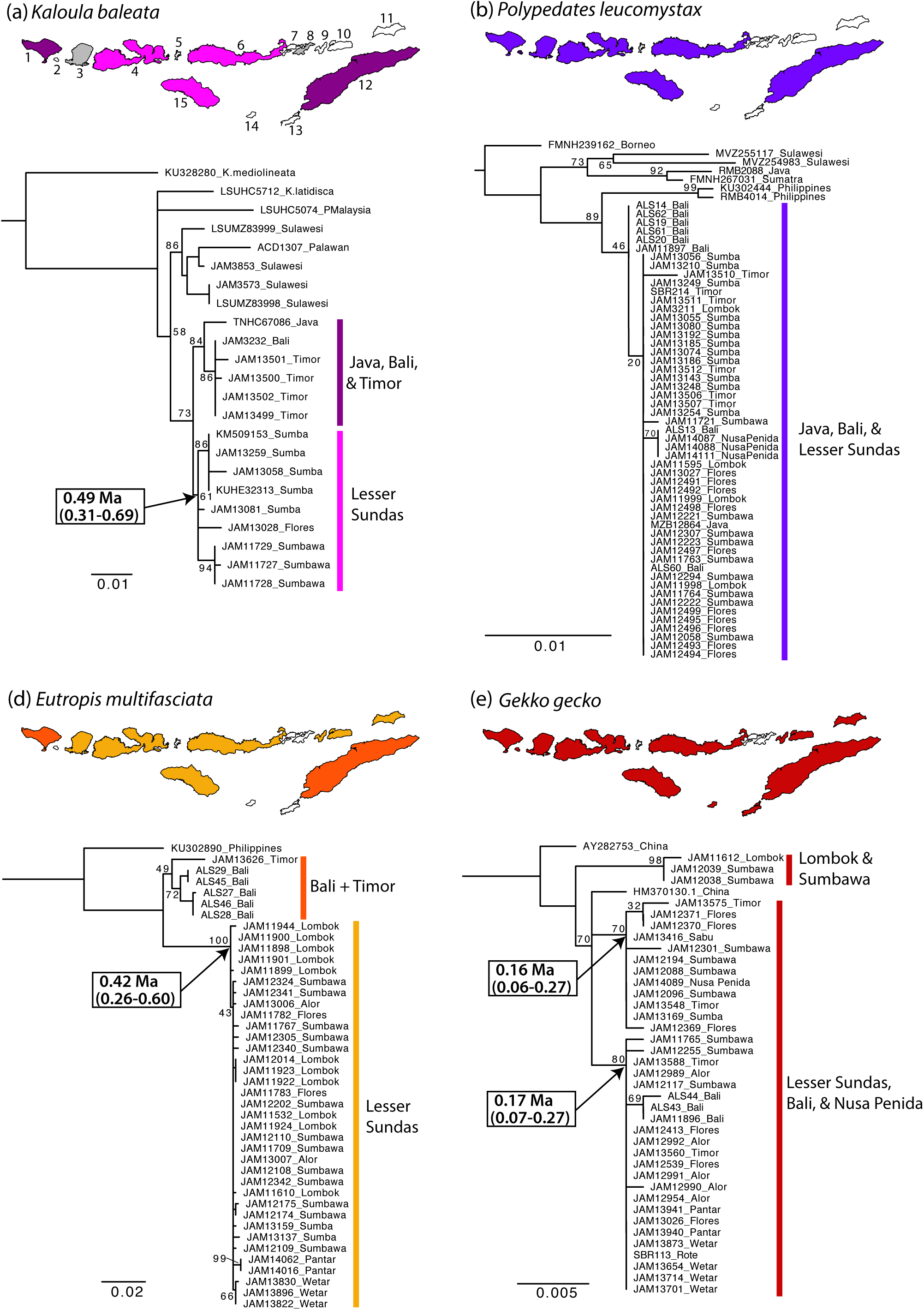
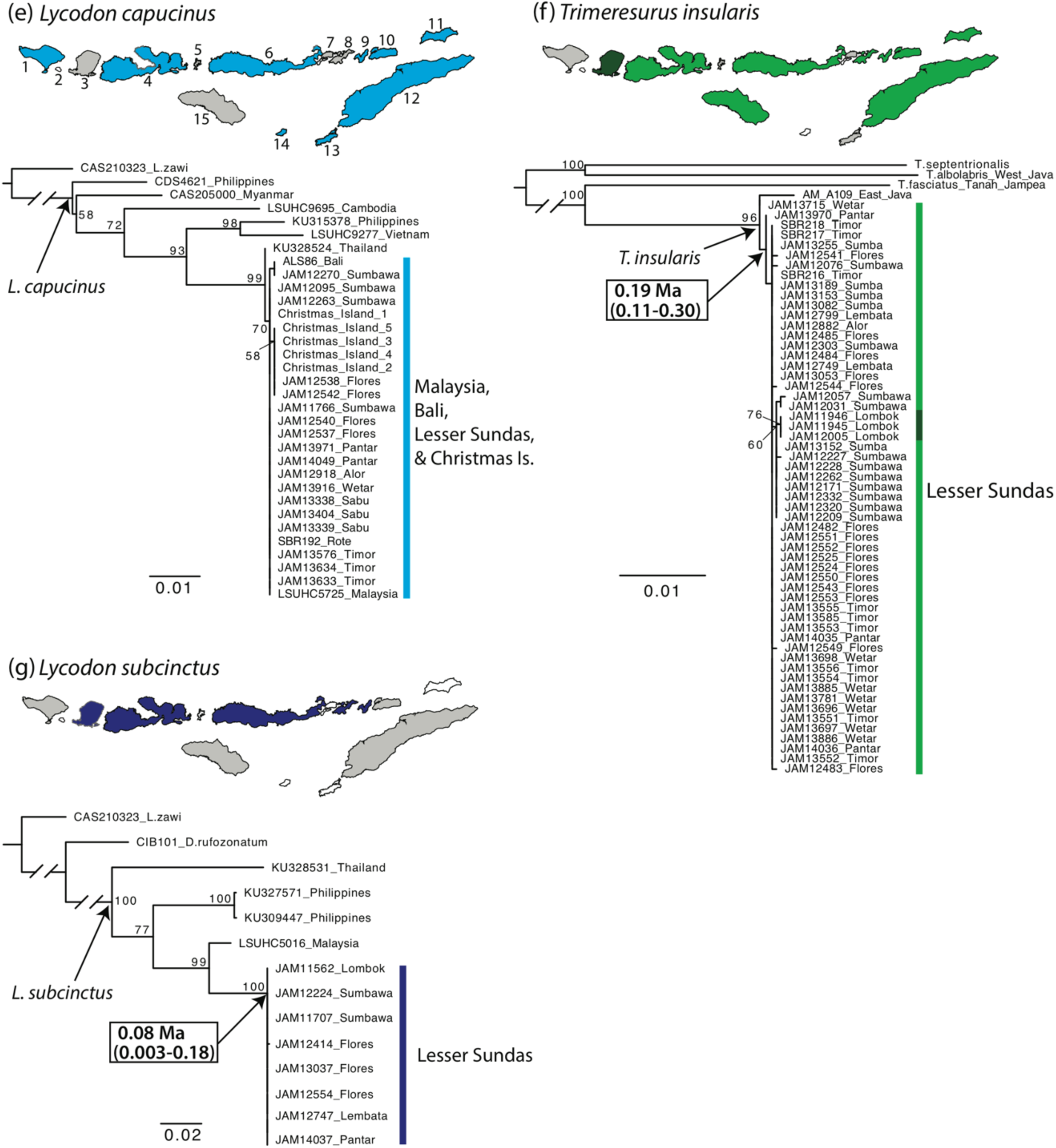
A-G. Maximum Likelihood phylogenies and represented islands. Each box contains a map of the Lesser Sundas region (including Bali and Nusa Penida) with sampled islands color filled. Unsampled islands where the species is known to occur are shaded light grey. Numbers at nodes represent bootstrap support. Text in boxes represents most recent common ancestor (MRCA) age estimates for relevant lineages. The numbers alongside islands in box A and E correspond to relevant islands: 1-Bali, 2-Nusa Penida, 3-Lombok, 4-Sumbawa, 5-Komodo, 6-Flores, 7-Adonara, 8-Lembata, 9-Pantar, 10-Alor, 11-Wetar, 12-Timor, 13-Rote, 14-Sabu, 15-Sumba.

Within *P. leucomystax*, the Lesser Sundas samples are most closely related to samples from Java, Bali, and Nusa Penida (Fig. 3b). A single common haplotype is shared across every island sampled within the Lesser Sundas as well as Java and Bali.

Within *E. multifasciata*, our single sample from Timor is sister to a Bali clade, though still highly divergent. This Bali/Timor clade is sister to all remaining samples forming a well-supported Lesser Sundas clade (bootstrap proportion=100) (Fig. 3c). These two haplotype groups are differentiated by ∼4% sequence divergence at the *ND4* gene. Within the Lesser Sundas, lizards from Wetar and Pantar each form monopheletic groups, though only supported by one informative mutation for each clade. A single common haplotype is found on Lombok, Sumbawa, Flores, and Alor.

Samples of *G. gecko* from the Lesser Sundas are found in three separate clades each comprised of a collection of samples representing multiple overlapping localities. For example, samples from Sumbawa are present in each of the three clades. One of the two samples from China is nested in amongst the three Lesser Sundas clades and a second sample from China is sister to all other samples (Fig. 3d). The only monophyletic island is Bali, united by a single shared mutation. One common haplotype is found on Nusa Penida, Sumbawa, Sumba, Sabu, and Timor, while a second common haplotype is found on Sumbawa, Flores, Pantar, Alor, Wetar, Timor, and Rote.

*Lycodon capucinus* samples from the Lesser Sundas contain a total of three haplotypes distinguished by two parsimony informative substitutions. The most common haplotype was found in samples from Malaysia, Sumbawa, Flores, Pantar, Alor, Wetar, Timor, Rote, Sabu, and Christmas Island (Fig. 3e). A second haplotype occurs on Christmas Island and Flores, and a third haplotype occurs on Bali and Sumbawa.

Within *L. subcinctus*, a total of two haplotypes were recovered from the eight newly sequenced samples with only one singleton mutation found in a Flores sample (Fig. 3g). The common haplotype was found in snakes from Lombok, Sumbawa, Flores, Lembata, and Pantar. The Lesser Sundas samples form a strongly supported clade relative to samples from Malaysia, Thailand, and the Philippines, but we lack samples from adjacent Java or Bali.

A total of 11 haplotypes were recovered from the 57 *T. insularis* samples with only three parsimony informative sites present. The most common haplotype, belonging to 40/57 Lesser Sunda samples, is found on every island except Lombok. Lombok is the only monophyletic island population and is differentiated by a single unique mutation (Fig. 3f). A *T. insularis* sample from eastern Java is sister to all Lesser Sundas samples. *Trimeresurus fasciatus* from Tanahjampea Island (123 km north of Flores) was found to be the sister taxon of *T. insularis*.

### 3.2 Summary statistics

Within the Lesser Sundas samples, genetic variability ranged from a low of one variable site and no parsimony informative sites (*L. subcinctus*) to a high of 54 variable sites with 10 parsimony informative sites (*E. multifasciata* “Lesser Sundas” clade + Timor sample) (Table 1). However, when *E. multifasciata* samples from the Lesser Sundas without Timor are considered there are only 23 variable sites, eight of which are parsimony informative. Haplotype diversity within Lesser Sundas samples ranged from 0.10 (*P. leucomystax*) to 0.91 (*E. multifasciata* “Lesser Sundas” clade) (Table 2). Nucleotide diversities within Lesser Sundas samples were all low and ranged from a low of 0.0002 (*P. leucomystax* and *L. subcinctus*) to a high of 0.0044 (*G. gecko*) (Table 2). Tajima’s D statistics calculated from Lesser Sundas samples were negative for all species, ranging from −0.58 (*K. baleata*) to −2.20 (statistically significant value for *E. multifasciata* “Lesser Sundas” clade). Fu’s *Fs* statistics were negative for all Lesser Sundas samples, ranging from −0.18 (*L. subcinctus*) to −16.52 (*E. multifasciata* “Lesser Sundas” clade), while the clades of *E. multifasciata* and *K. baleata* containing samples from Timor and Bali had positive values (2.55 and 1.02, respectively).

**TABLE 2.** Mitochondrial marker summary statistics and divergence times. h= haplotype diversity, π= nucleotide diversity; MRCA= most recent common ancestor; Ma= millions of years ago; *= statistically significant.

| Species | h | $\pi$ | Tajima's *D* | Fu's *F<sub>s</sub>* |
| --- | --- | --- | --- | --- |
| *Kaloula baleata* | - | - | - | - |
| - "Java/Bali/Timor" clade | 0.60 | 0.0031 | -1.37 (P>0.10) | 1.02 |
| - "Lesser Sundas" clade | 0.83 | 0.0037 | -0.58 (P>0.10) | -0.85 |
| *Polypedates leucomystax* | 0.10 | 0.0002 | -1.49 (P>0.10) | -2.66 |
| *Eutropis multifasciata* | - | - | - | - |
| - "Bali/Timor" clade | 0.87 | 0.0117 | -0.39 (P>0.10) | 2.55 |
| - "Lesser Sundas" clade | 0.91 | 0.0024 | -2.20 (P<0.01)* | -16.52 |
| *Gekko gecko* | 0.72 | 0.0044 | -0.77 (P>0.10) | -0.81 |
| *Lycodon capucinus* | 0.29 | 0.0003 | -1.12 (P>0.10) | -1.15 |
| *Lycodon subcinctus* | 0.25 | 0.0002 | -1.05 (P>0.10) | -0.18 |
| *Trimeresurus insularis* | 0.47 | 0.0006 | -1.36 (P>0.10) | -5.19 |

### 3.3 Ages of Lesser Sundas clades

The ages of the most recent common ancestors (MRCA) of each Lesser Sundas clade fell within the late Pleistocene (even considering 95% confidence intervals) with a range of 80,000-490,000 years (Figure 3). For the frogs, MRCA age was estimated for the Lesser Sundas clade (not including Timor) of *K. baleata* (0.49 Ma; 95% CI 0.31-0.69 Ma), and no estimate is given for *P. leucomystax* due to the nested Sundaland samples from Java, Bali, and Nusa Penida. Among lizard species, MRCA ages were estimated for the Lesser Sundas lineage (not including Timor) of *E. multifasciata* (0.42 Ma; 95% CI 0.26-0.60 Ma), and for two lineages of *G. gecko* that contain mostly Lesser Sundas samples: 1) a lineage containing samples from Sumbawa, Flores, Sumba, Sabu, Timor, and Nusa Penida (0.16 Ma; 95% CI 0.06-0.27 Ma), and 2) a lineage containing samples from Sumbawa, Flores, Pantar, Alor, Wetar, Timor, Rote, and Bali (0.17 Ma; 95% CI 0.07-0.27 Ma). Within snakes, MRCA ages were estimated for Lesser Sundas lineages of *L. subcinctus* (0.08 Ma; 95% CI 0.003-0.18 Ma) and *T. insularis* (0.19 Ma; 95% CI 0.11-0.30 Ma), and no estimate is given for *L. capucinus* because no monophyletic Lesser Sunda lineage exists.

## 4 DISCUSSION

### 4.1 General patterns

The mitochondrial markers we have screened are well-suited for rejecting human-mediated introductions since deep genetic structure would indicate that their occurrence in the region predated the arrival of humans. However, lack of structure does not necessarily mean that these species did not arrive via natural dispersal. We set out to determine if there was sufficiently deep genetic structure within these taxa to reject the hypothesis that the Lesser Sundas populations are the result of human introductions. However, one potential complication is that independent introductions from structured populations outside of the Lesser Sundas could be misinterpreted as structure within the Lesser Sundas. Adding to this challenge is the possibility that some taxa may be composed of both native and non-native populations.

If any of these species had been long-established in the Lesser Sundas, we would expect each island population, or at least sets of islands, to have unique mutations resulting in phylogeographic structure. The results of our mitochondrial phylogenetic estimates and summary statistics have shown that each of the seven focal species have little to moderate genetic differentiation and little to no phylogeographic structure across the oceanic islands of the Lesser Sundas. This pattern could be created by different scenarios, four of which are considered here: 1) natural introduction with prolific recent natural dispersals between islands, 2) natural introduction with human-mediated dispersal between islands, 3) a single human mediated introduction and human-mediated dispersal between islands, or 4) multiple human-mediated introductions and human-mediated dispersal between islands.

While our MRCA ages are only rough estimates, they do suggest that some species may have occurred within the Lesser Sundas on the order of hundreds of thousands of years, and the negative Tajima’s D values estimated for Lesser Sundas samples within each species are consistent with recent population expansions. Given that each of these species is known to have occurred throughout the majority of their currenty known Lesser Sundas range for over 100 years, any human mediated introductions into the archipelago or specific islands would have occurred before the 20^th^ century (Table S2). While a primary goal of this study was to search for evidence of a natural invasion for these taxa, we can reject human-mediated introduction to the archipelago for only *K. baleata, E. multifasciata*, and *T. insularis*. Even for these taxa, the general absence of phylogeographic structure within the Lesser Sundas indicates that we cannot reject recent human-mediated inter-island dispersal.

### 4.2 Candidates for natural introduction with possible human-mediated dispersal

The white-lipped island pitviper has very low genetic diversity with a common haplotype found on every island except Lombok. However, *T. insularis* does contain some phylogeographical structure with Lombok island samples forming a monophyletic group. Although the topology of our tree places samples from Wetar plus Pantar as basal within the Lesser Sundas, support for this branching pattern is very weak and it is more appropriate to interpret the tree as a large polytomy or star phylogeny. *Trimeresurus insularis* diverged from its sister species *T. fasciatus*, which is found on Tanahjampea Island to the North of Flores, approximately 2.4 Ma. It is unclear if the ancestor of these two species dispersed from Tanahjampea to the Lesser Sundas, or from the Lesser Sundas to Tanahjampea, but the prevailing north-to-south oceanic currents in the Flores Sea suggest that a dispersal from Tanahjampea to the Lesser Sundas is more likely (Gordon, 2005). If this is the case, then East Java may have been colonized by *T. insularis* from the Lesser Sundas. While it appears that this species has naturally occurred in the archipelago for some time, we note that both the high densities of *T. insularis* and the movement of large quantities of agricultural goods by boat could potentially transport these pitvipers between islands.

Both *Kaloula baleata* and *Eutropis multifasciata* have moderate phylogeographic structure and genetic diversity within the Lesser Sundas suggesting that they have inhabited the archipelago for at least thousands of years. Interestingly, for both species, our data indicate that samples from Timor are more closely related to samples from Bali (and Java in the case of *Kaloula*) than to samples representing the remainder of the Lesser Sundas. An important difference between these species, however, is that the *K. baleata* sample from Bali had an identical haplotype to some Timor samples, whereas the *E. multifasciata* sample from Timor is clearly divergent and quite genetically distinct from samples from Bali. This suggests that *K. baleata* likely arrived on Timor via human-mediated introduction from Bali, whereas *E. multifaciata* on Timor may very well represent a separate natural introduction from Bali or elsewhere. Regarding the non-Timor populations of *Kaloula* in the Lesser Sundas, our limited data set exhibits phylogeographic structure suggesting that these frogs are naturally-occurring on the islands of Sumbawa, Sumba, Flores, and likely others as well.

*Eutropis multifasciata* populations on Bali, Pantar, and Wetar are each monophyletic suggesting isolation on those islands, whereas Lombok, Sumbawa, Flores, Sumba, and Alor share either a common haplotype or very closely related haplotypes suggesting either recent or continued movement between those islands. *Eutropis multifasciata* has likely occurred in the Lesser Sundas for a significant amount of time but it is unclear if they have been moved between islands by people or if they are such prolific natural dispersers that they remain genetically unstructured. Future sampling of both genes and localities throughout their range will certainly shed light on this unresolved question.

### 4.3 Candidates for human-mediated introduction and dispersal

There is low phylogeographic structure within *P. leucomystax* from the Lesser Sundas and one common haplotype is found throughout the Lesser Sundas, as well as on Java and Bali. The extremely low nucleotide diversity (0.0002) suggests a recent and rapid spread through the Lesser Sundas. Though not yet proven, it has long been suspected that *P. leucomystax* has been moved around much of Southeast Asia by people. Our findings support the hypotheses that *P. leucomystax* has either been recently introduced into and throughout the Lesser Sundas Archipelago or that it has naturally invaded the archipelago very recently followed by rapid colonization of nearly every major island.

Analyses of the snake fauna within the Lesser Sundas suggest that the archipelago’s long-standing isolation from other islands in the region have influenced both their species composition and levels of variation between island populations (How & Kitchener, 1997). However, we find that both species of *Lycodon* wolf snakes have very little genetic diversity, no phylogeographical structure, and negative Tajima’s D values, suggesting that they have recently expanded their ranges and population sizes as they colonized the islands. *Lycodon capucinus* from the Lesser Sundas are nearly genetically identical to those from the Malay Peninsula suggesting they were recently introduced by humans, and likely continue to be moved between islands. *Lycodon subcinctus* also exhibits minimal genetic diversity suggesting a recent and rapid spread through the archipelago and/or continued movement between islands. However, the lack of *L. subcinctus* sampling from the nearby Sunda Shelf islands of Java and Bali, as well as the monophyly of Lesser Sundas samples prevent us from ruling out a natural introduction.

### 4.4 Candidate for multiple human-mediated introductions and dispersals

Movement of reptiles between islands in the Lesser Sundas is certainly known for some species such as the geckos *Hemidactylus frenatus, H. platyurus*, and *Gehyra mutilata*, which are commonly seen on ferry boats travelling between islands (pers. obs. S. Reilly). This may be the case for *G. gecko* as well, which has more genetic diversity than the three snakes examined but no discernable phylogeographic structure. *ND4* sequence data for *G. gecko* outside the Lesser Sundas is sparse which prevents the inference of the source populations for the Lesser Sundas. *Gekko gecko* samples within the Lesser Sundas are rendered paraphyletic and nested within samples from China suggesting multiple human mediated introductions. The introduction of the Chinese turtle, *Mauremys reevesii*, to Timor is evidence of the long history of travel and movement of animals between China and Indonesia (Yuwono, 1998; Kaiser *et al.*, 2010). *Gekko gecko* are common in dense human settlements in the region and are known to prey on smaller geckos (such as *Hemidactylus, Lepidodactylus*, and *Gehyra*), which are common on boats travelling between islands (Aowphol *et al.*, 2006). Thus, it would not be surprising if *G. gecko* occasionally or routinely stows away on these boats and is thereby moved between islands.

### 4.5 Conservation implications

Plants and animals have been moved between islands and introduced to new islands, both purposefully and unintentionally, within the Indo-Australian archipelago for thousands of years (Heinsohn, 2003). Certainly, the introduction of some species results in greater ecological change than others, and special caution should be given to prevent the human-mediated spread of those species that could cause major damage such as toxic species (e.g. toads) or certain predators that can devastate naïve native species. For example, the nearby Christmas Island has been heavily impacted by the human-mediated introduction of *L. capucinus*, which has likely caused the extinction of four species of lizards endemic to the island (Smith *et al.*, 2012). The detection of human-mediated introductions of ecologically harmful species will depend on accurate surveys of each island’s fauna both in the past (to determine a baseline) and the present (to detect introductions), and as such we recommend that comprehensive faunal surveys of the Lesser Sundas continue.

## 5 CONCLUSIONS

The results from this study show that none of these seven taxa exhibit deep genetic structure as seen in old, naturally occurring taxa such as *Limnonectes* fanged frogs, *Draco* flying lizards, and *Sphenomorphus* forest skinks (Reilly, 2016; Reilly *et al.*, 2019). In light of these results, we cannot definitively reject the hypothesis of human-mediated introductions and/or movements throughout the archipelago for these seven species. For some of these species (*K. baleata, E. multifasciata, T. insularis*, and *L. subcinctus*), longer term natural introductions remain a viable possibility that requires further investigation, whereas for others (*P. leucomystax, G. gecko, L. capucinus*) natural introductions seem unlikely in light of these new results, what we know about well-documented introductions elsewhere (such as for *L. capucinus*), and the biology of the species (rampant overwater dispersal over a short temporal extent would be quite unexpected for amphibians). While we have not definitively answered the question of the mode of introduction for most of these taxa, our data clearly indicate that each of these taxa is a relatively recent arrival to the Lesser Sundas Archipelago. We have shed light on the biogeography of each of these taxa and set the stage for others to follow-up with more comprehensive sampling of localities and genes.

## ACKNOWLEDGEMENTS

We thank Umilaela Arifin, Gilang Ramadhan, Jerome Fuchs, Jim and Carol Patton, Amir Hamidy, Kristopher Harmon, Luke Bloch, and Sarah Hykin for their help with the field collection of specimens and tissues. We thank Lydia Smith and the Evolutionary Genetics Laboratory (EGL) at UC Berkeley for laboratory support, Carol Spencer for accessioning of specimens, and Vishruth Venkataraman, Stephanie Wong, Saachi Gupta, Amanda Radel, Chantelle Khambolja, and Jennifer Lara for help with molecular lab work. Funding was provided by the National Geographic Society and the National Science Foundation (#DEB-1258185 awarded to JAM). Fieldwork in Indonesia was carried out under research permits issued by LIPI and RISTEK, and UC Berkeley IACUC protocol #R279.

## SUPPLEMENTAL MATERIAL

**TABLE S1.** Dates of the first records of each species on major islands in the Lesser Sundas. Numbers in brackets correspond to the supplemental literature cited below.

| Species | Bali | Lombok | Sumbawa | Sumba | Komodo | Flores | Lembata | Alor | Wetar | Savu | Rote | Timor |
| --- | --- | --- | --- | --- | --- | --- | --- | --- | --- | --- | --- | --- |
| *Kaloula baleata* | 1927 [1] | 1927 [1] | 1927 [1] | 1897 [5] | 1927 [1] | 1927 [1] | 2011 [7] |  |  |  |  | 2009 [7] |
| *Polypedates leucomystax* | 1927 [1] | 1907 [6] | 1927 [1] | 1897 [5] |  | 1927 [1] |  |  |  |  |  | 1923 [3] |
| *Eutropis multifasciatus* | 1927 [1] | 1897 [5] | 1917 [2] |  | 1927 [1] | 1917 [2] |  | 1917 [2] | 1917 [2] |  |  | 1917 [2] |
| *Gekko gecko* | 1927 [1] | 1897 [5] | 1927 [1] | 1917 [2] | 1927 [1] | 1917 [2] |  | 1917 [2] |  | 1897 [5] | 1898 [4] | 1898 [4] |
| *Lycodon capucinus* | 1917 [2] |  | 1917 [2] | 1897 [5] | 1927 [1] | 1917 [2] | 1917 [2] | 1917 [2] | 1917 [2] | 1897 [5] | 1898 [4] | 1917 [2] |
| *Lycodon subcinctus* |  | 1897 [5] | 1917 [2] |  |  | 1958 [8] |  |  |  |  |  |  |
| *Trimeresurus insularis* |  | 1927 [1] | 1927 [1] |  | 1927 [1] | 1927 [1] |  |  |  |  | 1898 [4] | 1898 [4] |

**\*Supplemental Literature**
- 1- Mertens, R. (1930) Die Amphibien und Reptilien der Inseln Bali, Lombok, Sumbawa and Flores. *Abhandlungen Senckenbergischen Naturforschenden Gesellschaft*, 42, 115–344. - 2- de Rooij, N. (1917). *The Reptiles of the Indo-Australian Archipelago* (Vol. 1-2). EJ Brill. - 3- van Kampen, P. N. (1923). *The amphibia of the Indo-Australian Archipelago*. E. J. Brill, Ltd, Leiden, 304 pp. - 4- van Lidth de Jeude, T. W. (1895). Reptiles from Timor and the neighbouring islands. *Notes from the Leyden Museum*, 16, 119-127. - 5- Boulenger, G. A. (1897). LII.—A list of the reptiles and batrachians collected by Mr. Alfred Everett in Lombok, Flores, Sumba, and Savu, with descriptions of new species. *Journal of Natural History*, 19, 503-509. - 6- Barbour, T. (1912). A contribution to the zoogeography of the East Indian Islands. *Memoirs of the Museum of Comparative Zoology*, 44, 1-168. - 7- Kaiser, H., Carvalho, V. L., Ceballos, J., Freed, P., Heacox, S., Lester, B., Richards, S. J., Trainor, C. R., Sanchez, C. & O'Shea, M. (2011). The herpetofauna of Timor-Leste: a first report. *ZooKeys*, (109), 19. - 8- van Hoesel, J. P. K. (1959) *Ophidia Javanica*. Museum Zoologicum Bogoriense, Bogor, Indonesia. 144 pp.

**TABLE S2.** Museum numbers and locality information for samples used in this study.

| Museum # | Field Catalog # | Species | Island | Latitude | Longitude |
| --- | --- | --- | --- | --- | --- |
| MVZ274408 | ALS27 | *Eutropis multifasciata* | Bali | -8.43810 | 115.40172 |
| MVZ274409 | ALS28 | *Eutropis multifasciata* | Bali | -8.43810 | 115.40172 |
| MVZ274410 | ALS29 | *Eutropis multifasciata* | Bali | -8.43810 | 115.40172 |
| MVZ274411 | ALS45 | *Eutropis multifasciata* | Bali | -8.43810 | 115.40172 |
| MVZ274412 | ALS46 | *Eutropis multifasciata* | Bali | -8.43810 | 115.40172 |
| MVZ291606 | JAM11532 | *Eutropis multifasciata* | Lombok | -8.52854 | 116.28206 |
| MVZ291607 | JAM11610 | *Eutropis multifasciata* | Lombok | -8.53159 | 116.39878 |
| MVZ291608 | JAM11709 | *Eutropis multifasciata* | Sumbawa | -8.41016 | 117.21491 |
| MVZ291609 | JAM11767 | *Eutropis multifasciata* | Sumbawa | -8.73561 | 117.73106 |
| MVZ291610 | JAM11782 | *Eutropis multifasciata* | Flores | -8.54530 | 119.91059 |
| MVZ291611 | JAM11783 | *Eutropis multifasciata* | Flores | -8.54530 | 119.91059 |
| MVZ291612 | JAM11898 | *Eutropis multifasciata* | Lombok | -8.53912 | 116.53959 |
| MVZ291613 | JAM11899 | *Eutropis multifasciata* | Lombok | -8.53912 | 116.53959 |
| MVZ291614 | JAM11900 | *Eutropis multifasciata* | Lombok | -8.53912 | 116.53959 |
| MVZ291615 | JAM11901 | *Eutropis multifasciata* | Lombok | -8.53912 | 116.53959 |
| MVZ291616 | JAM11922 | *Eutropis multifasciata* | Lombok | -8.36115 | 116.52200 |
| MVZ291617 | JAM11923 | *Eutropis multifasciata* | Lombok | -8.36115 | 116.52200 |
| MVZ291618 | JAM11924 | *Eutropis multifasciata* | Lombok | -8.36115 | 116.52200 |
| MVZ291619 | JAM11944 | *Eutropis multifasciata* | Lombok | -8.36115 | 116.52200 |
| MVZ291620 | JAM12014 | *Eutropis multifasciata* | Lombok | -8.36115 | 116.52000 |
| MVZ291621 | JAM12108 | *Eutropis multifasciata* | Sumbawa | -8.91228 | 116.75224 |
| MVZ291622 | JAM12109 | *Eutropis multifasciata* | Sumbawa | -8.51717 | 118.31950 |
| MVZ291623 | JAM12110 | *Eutropis multifasciata* | Sumbawa | -8.51717 | 118.31950 |
| MVZ291624 | JAM12174 | *Eutropis multifasciata* | Sumbawa | -8.54700 | 118.46947 |
| MVZ291625 | JAM12175 | *Eutropis multifasciata* | Sumbawa | -8.54700 | 118.46947 |
| MVZ291626 | JAM12202 | *Eutropis multifasciata* | Sumbawa | -8.67775 | 118.68163 |
| MVZ291627 | JAM12305 | *Eutropis multifasciata* | Sumbawa | -8.76643 | 118.60496 |
| MVZ291628 | JAM12324 | *Eutropis multifasciata* | Sumbawa | -8.76296 | 118.61102 |
| MVZ291629 | JAM12340 | *Eutropis multifasciata* | Sumbawa | -8.59961 | 119.01229 |
| MVZ291630 | JAM12341 | *Eutropis multifasciata* | Sumbawa | -8.59961 | 119.01229 |
| MVZ291631 | JAM12342 | *Eutropis multifasciata* | Sumbawa | -8.59961 | 119.01229 |
| MVZ291632 | JAM13006 | *Eutropis multifasciata* | Alor | -8.17922 | 124.55897 |
| MVZ291633 | JAM13007 | *Eutropis multifasciata* | Alor | -8.17922 | 124.55897 |
| MVZ291634 | JAM13137 | *Eutropis multifasciata* | Sumba | -10.02102 | 120.05795 |
| MVZ291635 | JAM13159 | *Eutropis multifasciata* | Sumba | -10.02102 | 120.05795 |
| MVZ291636 | JAM13822 | *Eutropis multifasciata* | Wetar | -7.92789 | 126.40831 |
| MVZ291637 | JAM13830 | *Eutropis multifasciata* | Wetar | -7.92235 | 126.40569 |
| MVZ291638 | JAM13896 | *Eutropis multifasciata* | Wetar | -7.91965 | 126.39785 |
| MVZ291639 | JAM14016 | *Eutropis multifasciata* | Pantar | -8.35268 | 124.25156 |
| MVZ291640 | JAM14062 | *Eutropis multifasciata* | Pantar | -8.35392 | 124.25345 |
| MVZ291641 | JAM13626 | *Eutropis multifasciata* | Timor | -10.03848 | 123.93333 |
| MVZ274447 | ALS43 | *Gekko gekko* | Bali | -8.43810 | 115.40172 |
| MVZ274448 | ALS44 | *Gekko gekko* | Bali | -8.43810 | 115.40172 |
| MVZ291642 | SBR113 | *Gekko gekko* | Rote | -10.88656 | 122.82613 |

**TABLE S2 cont.**
| **Museum #** | **Field Catalog #** | **Species** | **Island** | **Latitude** | **Longitude** |
| --- | --- | --- | --- | --- | --- |
| MVZ291643 | JAM11612 | *Gekko gekko* | Lombok | -8.58987 | 116.12589 |
| MVZ291644 | JAM11765 | *Gekko gekko* | Sumbawa | -8.67102 | 118.23570 |
| MVZ291645 | JAM11896 | *Gekko gekko* | Bali | -8.81452 | 115.10137 |
| MVZ291646 | JAM12038 | *Gekko gekko* | Sumbawa | -9.02763 | 116.81893 |
| MVZ291647 | JAM12039 | *Gekko gekko* | Sumbawa | -9.02763 | 116.81893 |
| MVZ291648 | JAM12088 | *Gekko gekko* | Sumbawa | -8.76137 | 118.00056 |
| MVZ291649 | JAM12096 | *Gekko gekko* | Sumbawa | -8.60250 | 118.28671 |
| MVZ291650 | JAM12117 | *Gekko gekko* | Sumbawa | -8.43721 | 118.30004 |
| MVZ291651 | JAM12194 | *Gekko gekko* | Sumbawa | -8.51728 | 118.31947 |
| MVZ291652 | JAM12255 | *Gekko gekko* | Sumbawa | -8.49715 | 118.80463 |
| MVZ291653 | JAM12301 | *Gekko gekko* | Sumbawa | -8.50065 | 119.03625 |
| MVZ291654 | JAM12369 | *Gekko gekko* | Flores | -8.78828 | 121.67159 |
| MVZ291655 | JAM12370 | *Gekko gekko* | Flores | -8.78828 | 121.67159 |
| MVZ291656 | JAM12371 | *Gekko gekko* | Flores | -8.78828 | 121.67159 |
| MVZ291657 | JAM12413 | *Gekko gekko* | Flores | -8.54537 | 119.91075 |
| MVZ291658 | JAM12539 | *Gekko gekko* | Flores | -8.61050 | 122.39379 |
| MVZ291659 | JAM12954 | *Gekko gekko* | Alor | -8.16476 | 124.74838 |
| MVZ291660 | JAM12989 | *Gekko gekko* | Alor | -8.18862 | 124.75208 |
| MVZ291661 | JAM12990 | *Gekko gekko* | Alor | -8.19025 | 124.75142 |
| MVZ291662 | JAM12991 | *Gekko gekko* | Alor | -8.19025 | 124.75142 |
| MVZ291663 | JAM12992 | *Gekko gekko* | Alor | -8.19025 | 124.75142 |
| MVZ291664 | JAM13026 | *Gekko gekko* | Flores | -8.21648 | 122.97288 |
| MVZ291665 | JAM13169 | *Gekko gekko* | Sumba | -10.01944 | 120.05637 |
| MVZ291666 | JAM13416 | *Gekko gekko* | Sabu | -10.50747 | 121.99487 |
| MVZ291667 | JAM13548 | *Gekko gekko* | Timor | -10.02225 | 123.86884 |
| MVZ291668 | JAM13560 | *Gekko gekko* | Timor | -10.10636 | 123.71033 |
| MVZ291669 | JAM13575 | *Gekko gekko* | Timor | -10.02339 | 123.86659 |
| MVZ291670 | JAM13588 | *Gekko gekko* | Timor | -10.03903 | 123.93174 |
| MVZ291671 | JAM13654 | *Gekko gekko* | Wetar | -7.93010 | 126.41180 |
| MVZ291672 | JAM13701 | *Gekko gekko* | Wetar | -7.92621 | 126.40793 |
| MVZ291673 | JAM13714 | *Gekko gekko* | Wetar | -7.92686 | 126.41972 |
| MVZ291674 | JAM13873 | *Gekko gekko* | Wetar | -7.92237 | 126.40568 |
| MVZ291675 | JAM13940 | *Gekko gekko* | Pantar | -8.35555 | 124.25712 |
| MVZ291676 | JAM13941 | *Gekko gekko* | Pantar | -8.35555 | 124.25712 |
| MVZ291677 | JAM14089 | *Gekko gekko* | Nusa Penida | -8.68072 | 115.48843 |
| MVZ291190 | JAM11945 | *Trimeresurus insularis* | Lombok | -8.25729 | 116.44359 |
| MVZ291191 | JAM11946 | *Trimeresurus insularis* | Lombok | -8.30545 | 116.40701 |
| MVZ291192 | JAM12005 | *Trimeresurus insularis* | Lombok | -8.30634 | 116.40496 |
| MVZ291193 | JAM12031 | *Trimeresurus insularis* | Sumbawa | -9.00542 | 116.75720 |
| MVZ291194 | JAM12057 | *Trimeresurus insularis* | Sumbawa | -9.05051 | 116.86373 |
| MVZ291195 | JAM12076 | *Trimeresurus insularis* | Sumbawa | -9.00639 | 116.76723 |
| MVZ291196 | JAM12171 | *Trimeresurus insularis* | Sumbawa | -8.66990 | 118.43940 |
| MVZ291197 | JAM12209 | *Trimeresurus insularis* | Sumbawa | -8.74106 | 118.60437 |
| MVZ291198 | JAM12227 | *Trimeresurus insularis* | Sumbawa | -8.71271 | 118.62209 |

TABLE S2 cont.
| Museum # | Field Catalog # | Species | Island | Latitude | Longitude |
| --- | --- | --- | --- | --- | --- |
| MVZ291199 | JAM12228 | *Trimeresurus insularis* | Sumbawa | -8.54886 | 118.40537 |
| MVZ291200 | JAM12262 | *Trimeresurus insularis* | Sumbawa | -8.65415 | 118.09328 |
| MVZ291201 | JAM12303 | *Trimeresurus insularis* | Sumbawa | -8.57221 | 118.87612 |
| MVZ291202 | JAM12320 | *Trimeresurus insularis* | Sumbawa | -8.76643 | 118.60496 |
| MVZ291203 | JAM12332 | *Trimeresurus insularis* | Sumbawa | -8.76300 | 118.60707 |
| MVZ291204 | JAM12482 | *Trimeresurus insularis* | Flores | -8.68687 | 121.03889 |
| MVZ291205 | JAM12483 | *Trimeresurus insularis* | Flores | -8.68687 | 121.03889 |
| MVZ291206 | JAM12484 | *Trimeresurus insularis* | Flores | -8.68687 | 121.03889 |
| MVZ291207 | JAM12485 | *Trimeresurus insularis* | Flores | -8.68687 | 121.03889 |
| MVZ291208 | JAM12524 | *Trimeresurus insularis* | Flores | -8.75954 | 121.70090 |
| MVZ291209 | JAM12525 | *Trimeresurus insularis* | Flores | -8.75954 | 121.70090 |
| MVZ291210 | JAM12541 | *Trimeresurus insularis* | Flores | -8.54421 | 122.51745 |
| MVZ291211 | JAM12543 | *Trimeresurus insularis* | Flores | -8.49705 | 122.54664 |
| MVZ291212 | JAM12544 | *Trimeresurus insularis* | Flores | -8.41902 | 122.81760 |
| MVZ291213 | JAM12549 | *Trimeresurus insularis* | Flores | -8.48070 | 122.58190 |
| MVZ291214 | JAM12550 | *Trimeresurus insularis* | Flores | -8.48936 | 122.60113 |
| MVZ291215 | JAM12551 | *Trimeresurus insularis* | Flores | -8.49313 | 122.60400 |
| MVZ291216 | JAM12552 | *Trimeresurus insularis* | Flores | -8.50030 | 122.76844 |
| MVZ291217 | JAM12553 | *Trimeresurus insularis* | Flores | -8.41956 | 122.81806 |
| MVZ291218 | JAM12749 | *Trimeresurus insularis* | Lembata | -8.38181 | 123.40195 |
| MVZ291219 | JAM12799 | *Trimeresurus insularis* | Lembata | -8.38181 | 123.40195 |
| MVZ291220 | JAM12882 | *Trimeresurus insularis* | Alor | -8.18880 | 124.69309 |
| MVZ291221 | JAM13053 | *Trimeresurus insularis* | Flores | -8.68277 | 122.21175 |
| MVZ291222 | JAM13082 | *Trimeresurus insularis* | Sumba | -10.01906 | 120.05923 |
| MVZ291223 | JAM13152 | *Trimeresurus insularis* | Sumba | -10.02426 | 120.05718 |
| MVZ291224 | JAM13153 | *Trimeresurus insularis* | Sumba | -10.02426 | 120.05718 |
| MVZ291227 | JAM13189 | *Trimeresurus insularis* | Sumba | -10.02012 | 120.05795 |
| MVZ291228 | JAM13255 | *Trimeresurus insularis* | Sumba | -10.02193 | 120.05812 |
| MVZ291229 | JAM13551 | *Trimeresurus insularis* | Timor | -10.02225 | 123.86884 |
| MVZ291230 | JAM13552 | *Trimeresurus insularis* | Timor | -10.02225 | 123.86884 |
| MVZ291231 | JAM13553 | *Trimeresurus insularis* | Timor | -10.02225 | 123.86884 |
| MVZ291232 | JAM13554 | *Trimeresurus insularis* | Timor | -10.02225 | 123.86884 |
| MVZ291233 | JAM13555 | *Trimeresurus insularis* | Timor | -10.02225 | 123.86884 |
| MVZ291234 | JAM13556 | *Trimeresurus insularis* | Timor | -10.02225 | 123.86884 |
| MVZ291235 | JAM13597 | *Trimeresurus insularis* | Timor | -10.03903 | 123.93174 |
| MVZ291236 | JAM13696 | *Trimeresurus insularis* | Wetar | -7.92847 | 126.40781 |
| MVZ291237 | JAM13697 | *Trimeresurus insularis* | Wetar | -7.92769 | 126.40819 |
| MVZ291238 | JAM13698 | *Trimeresurus insularis* | Wetar | -7.92981 | 126.40920 |
| MVZ291239 | JAM13715 | *Trimeresurus insularis* | Wetar | -7.92686 | 126.41972 |
| MVZ291241 | JAM13781 | *Trimeresurus insularis* | Wetar | -7.92599 | 126.40797 |
| MVZ291244 | JAM13885 | *Trimeresurus insularis* | Wetar | -7.92853 | 126.40974 |
| MVZ291245 | JAM13886 | *Trimeresurus insularis* | Wetar | -7.92225 | 126.40114 |
| MVZ291246 | JAM13970 | *Trimeresurus insularis* | Pantar | -8.35124 | 124.25074 |
| MVZ291247 | JAM14035 | *Trimeresurus insularis* | Pantar | -8.35392 | 124.25347 |

TABLE S2 cont.
| Museum # | Field Catalog # | Species | Island | Latitude | Longitude |
| --- | --- | --- | --- | --- | --- |
| MVZ291248 | JAM14036 | *Trimeresurus insularis* | Pantar | -8.35392 | 124.25347 |
| MVZ291249 | SBR216 | *Trimeresurus insularis* | Timor | -9.82180 | 124.31034 |
| MVZ291250 | SBR217 | *Trimeresurus insularis* | Timor | -9.82180 | 124.31034 |
| MVZ291251 | SBR218 | *Trimeresurus insularis* | Timor | -9.82180 | 124.31034 |
| MVZ291678 | JAM11562 | *Lycodon subcinctus* | Lombok | -8.54447 | 116.39862 |
| MVZ291679 | JAM11707 | *Lycodon subcinctus* | Sumbawa | -8.57435 | 117.31339 |
| MVZ291680 | JAM12224 | *Lycodon subcinctus* | Sumbawa | -8.73993 | 118.60497 |
| MVZ291681 | JAM12414 | *Lycodon subcinctus* | Flores | -8.59420 | 119.96857 |
| MVZ291682 | JAM12554 | *Lycodon subcinctus* | Flores | -8.40184 | 122.84223 |
| MVZ291683 | JAM13037 | *Lycodon subcinctus* | Flores | -8.23836 | 122.97288 |
| MVZ291684 | JAM12747 | *Lycodon subcinctus* | Lembata | -8.38181 | 123.40195 |
| MVZ291685 | JAM14037 | *Lycodon subcinctus* | Pantar | -8.35392 | 124.25347 |
| MVZ273765 | ALS86 | *Lycodon capucinus* | Bali | -8.82545 | 115.09189 |
| MVZ291686 | JAM11766 | *Lycodon capucinus* | Sumbawa | -8.66042 | 118.08301 |
| MVZ291687 | JAM12095 | *Lycodon capucinus* | Sumbawa | -8.66243 | 118.26863 |
| MVZ291688 | JAM12263 | *Lycodon capucinus* | Sumbawa | -8.65627 | 118.08421 |
| MVZ291689 | JAM12270 | *Lycodon capucinus* | Sumbawa | -8.48579 | 118.78094 |
| MVZ291690 | JAM12537 | *Lycodon capucinus* | Flores | -8.61050 | 122.39379 |
| MVZ291691 | JAM12538 | *Lycodon capucinus* | Flores | -8.61061 | 122.39728 |
| MVZ291692 | JAM12540 | *Lycodon capucinus* | Flores | -8.57409 | 122.39379 |
| MVZ291693 | JAM12542 | *Lycodon capucinus* | Flores | -8.49705 | 122.54664 |
| MVZ291694 | JAM13971 | *Lycodon capucinus* | Pantar | -8.35268 | 124.25156 |
| MVZ291695 | JAM14049 | *Lycodon capucinus* | Pantar | -8.35591 | 124.25462 |
| MVZ291696 | JAM12918 | *Lycodon capucinus* | Alor | -8.16564 | 124.74774 |
| MVZ291697 | JAM13916 | *Lycodon capucinus* | Wetar | -7.92478 | 126.40736 |
| MVZ291698 | JAM13338 | *Lycodon capucinus* | Sabu | -10.51187 | 121.98785 |
| MVZ291699 | JAM13339 | *Lycodon capucinus* | Sabu | -10.51187 | 121.98785 |
| MVZ291700 | JAM13404 | *Lycodon capucinus* | Sabu | -10.50038 | 121.98094 |
| MVZ291701 | SBR192 | *Lycodon capucinus* | Rote | -10.74501 | 123.05447 |
| MVZ291702 | JAM13576 | *Lycodon capucinus* | Timor | -10.02945 | 123.36151 |
| MVZ291703 | JAM13633 | *Lycodon capucinus* | Timor | -10.26607 | 123.56586 |
| MVZ291704 | JAM13634 | *Lycodon capucinus* | Timor | -10.26607 | 123.56586 |
| - | Christmas Is. 1 | *Lycodon capucinus* | Christmas Island | -10.48775 | 105.64536 |
| - | Christmas Is. 2 | *Lycodon capucinus* | Christmas Island | -10.48775 | 105.64536 |
| - | Christmas Is. 3 | *Lycodon capucinus* | Christmas Island | -10.48775 | 105.64536 |
| - | Christmas Is. 4 | *Lycodon capucinus* | Christmas Island | -10.48775 | 105.64536 |
| - | Christmas Is. 5 | *Lycodon capucinus* | Christmas Island | -10.48775 | 105.64536 |
| MVZ291705 | JAM11727 | *Kaloula baleata* | Sumbawa | -8.72371 | 117.38319 |
| MVZ291706 | JAM11728 | *Kaloula baleata* | Sumbawa | -8.72371 | 117.38319 |
| MVZ291707 | JAM11729 | *Kaloula baleata* | Sumbawa | -8.72371 | 117.38319 |
| MVZ291708 | JAM13028 | *Kaloula baleata* | Flores | -8.21648 | 122.97288 |
| MVZ291709 | JAM13058 | *Kaloula baleata* | Sumba | -9.82975 | 119.92864 |
| MVZ291710 | JAM13081 | *Kaloula baleata* | Sumba | -10.02102 | 120.05795 |
| MVZ291711 | JAM13259 | *Kaloula baleata* | Sumba | -10.01994 | 120.05637 |

TABLE S2 cont.
| Museum # | Field Catalog # | Species | Island | Latitude | Longitude |
| --- | --- | --- | --- | --- | --- |
| MVZ291712 | JAM13499 | *Kaloula baleata* | Timor | -10.24424 | 123.65848 |
| MVZ291713 | JAM13500 | *Kaloula baleata* | Timor | -10.24424 | 123.65848 |
| MVZ291714 | JAM13501 | *Kaloula baleata* | Timor | -10.24424 | 123.65848 |
| MVZ291715 | JAM13502 | *Kaloula baleata* | Timor | -10.24424 | 123.65848 |
| MVZ273785 | ALS13 | *Polypedates leucomystax* | Bali | -8.82545 | 115.09189 |
| MVZ273786 | ALS14 | *Polypedates leucomystax* | Bali | -8.82545 | 115.09189 |
| MVZ273787 | ALS19 | *Polypedates leucomystax* | Bali | -8.82545 | 115.09189 |
| MVZ273788 | ALS20 | *Polypedates leucomystax* | Bali | -8.82545 | 115.09189 |
| MVZ273781 | ALS60 | *Polypedates leucomystax* | Bali | -8.43810 | 115.40172 |
| MVZ273782 | ALS61 | *Polypedates leucomystax* | Bali | -8.43810 | 115.40172 |
| MVZ273783 | ALS62 | *Polypedates leucomystax* | Bali | -8.43810 | 115.40172 |
| MVZ291716 | SBR214 | *Polypedates leucomystax* | Timor | -9.82180 | 124.31039 |
| MVZ291717 | JAM11595 | *Polypedates leucomystax* | Lombok | -8.53159 | 116.39878 |
| MVZ291718 | JAM11721 | *Polypedates leucomystax* | Sumbawa | -8.75751 | 117.34994 |
| MVZ291719 | JAM11763 | *Polypedates leucomystax* | Sumbawa | -8.52740 | 118.31392 |
| MVZ291720 | JAM11764 | *Polypedates leucomystax* | Sumbawa | -8.52740 | 118.31392 |
| MVZ291721 | JAM11897 | *Polypedates leucomystax* | Bali | -8.81452 | 115.10137 |
| MVZ291722 | JAM11998 | *Polypedates leucomystax* | Lombok | -8.30782 | 116.40483 |
| MVZ291723 | JAM11999 | *Polypedates leucomystax* | Lombok | -8.30782 | 116.40483 |
| MVZ291724 | JAM12221 | *Polypedates leucomystax* | Sumbawa | -8.73993 | 118.60497 |
| MVZ291725 | JAM12222 | *Polypedates leucomystax* | Sumbawa | -8.73993 | 118.60497 |
| MVZ291726 | JAM12223 | *Polypedates leucomystax* | Sumbawa | -8.73993 | 118.60497 |
| MVZ291727 | JAM12294 | *Polypedates leucomystax* | Sumbawa | -8.50065 | 119.03625 |
| MVZ291728 | JAM12307 | *Polypedates leucomystax* | Sumbawa | -8.76643 | 118.60496 |
| MVZ291729 | JAM12491 | *Polypedates leucomystax* | Flores | -8.68687 | 121.03889 |
| MVZ291730 | JAM12492 | *Polypedates leucomystax* | Flores | -8.68687 | 121.03889 |
| MVZ291731 | JAM12493 | *Polypedates leucomystax* | Flores | -8.68687 | 121.03889 |
| MVZ291732 | JAM12494 | *Polypedates leucomystax* | Flores | -8.68687 | 121.03889 |
| MVZ291733 | JAM12495 | *Polypedates leucomystax* | Flores | -8.68687 | 121.03889 |
| MVZ291734 | JAM12496 | *Polypedates leucomystax* | Flores | -8.68687 | 121.03889 |
| MVZ291735 | JAM12497 | *Polypedates leucomystax* | Flores | -8.68687 | 121.03889 |
| MVZ291736 | JAM12498 | *Polypedates leucomystax* | Flores | -8.68687 | 121.03889 |
| MVZ291737 | JAM12499 | *Polypedates leucomystax* | Flores | -8.68687 | 121.03889 |
| MVZ291738 | JAM13027 | *Polypedates leucomystax* | Flores | -8.21648 | 122.97288 |
| MVZ291739 | JAM13055 | *Polypedates leucomystax* | Sumba | -9.82975 | 119.92864 |
| MVZ291740 | JAM13056 | *Polypedates leucomystax* | Sumba | -9.82975 | 119.92864 |
| MVZ291741 | JAM13079 | *Polypedates leucomystax* | Sumba | -10.02102 | 120.05795 |
| MVZ291742 | JAM13080 | *Polypedates leucomystax* | Sumba | -10.02102 | 120.05795 |
| MVZ291743 | JAM13143 | *Polypedates leucomystax* | Sumba | -10.02783 | 120.05765 |
| MVZ291744 | JAM13185 | *Polypedates leucomystax* | Sumba | -10.02323 | 120.06006 |
| MVZ291745 | JAM13186 | *Polypedates leucomystax* | Sumba | -10.02323 | 120.06006 |
| MVZ291746 | JAM13192 | *Polypedates leucomystax* | Sumba | -10.01994 | 120.05637 |
| MVZ291747 | JAM13210 | *Polypedates leucomystax* | Sumba | -10.02323 | 120.06006 |
| MVZ291748 | JAM13248 | *Polypedates leucomystax* | Sumba | -10.00728 | 120.04520 |

**TABLE S2 cont.**
| **Museum #** | **Field Catalog #** | **Species** | **Island** | **Latitude** | **Longitude** |
| --- | --- | --- | --- | --- | --- |
| MVZ291749 | JAM13249 | *Polypedates leucomystax* | Sumba | -10.01994 | 120.05637 |
| MVZ291750 | JAM13254 | *Polypedates leucomystax* | Sumba | -10.02314 | 120.06021 |
| MVZ291751 | JAM13506 | *Polypedates leucomystax* | Timor | -10.24424 | 123.65848 |
| MVZ291752 | JAM13507 | *Polypedates leucomystax* | Timor | -10.24424 | 123.65848 |
| MVZ291753 | JAM13510 | *Polypedates leucomystax* | Timor | -10.24424 | 123.65848 |
| MVZ291754 | JAM13511 | *Polypedates leucomystax* | Timor | -10.24424 | 123.65848 |
| MVZ291755 | JAM13512 | *Polypedates leucomystax* | Timor | -10.24424 | 123.65848 |
| MVZ291756 | JAM14087 | *Polypedates leucomystax* | Nusa Penida | -8.71780 | 115.52113 |
| MVZ291757 | JAM14088 | *Polypedates leucomystax* | Nusa Penida | -8.71780 | 115.52113 |
| MVZ291758 | JAM14111 | *Polypedates leucomystax* | Nusa Penida | -8.73340 | 115.52335 |

## REFERENCES

Aowphol, A., Thirakhupt, K., Nabhitabhata, J., & Voris, H. K. (2006). Foraging ecology of the Tokay gecko, Gekko gecko, in a residential area in Thailand. Amphibia-Reptilia, 27, 491–503. https://doi.org/10.1163/156853806778877121

Arevalo, E., Davis, S. K., & Sites Jr, J. W. (1994). Mitochondrial DNA sequence divergence and phylogenetic relationships among eight chromosome races of the Sceloporus grammicus complex (Phrynosomatidae) in central Mexico. Systematic Biology, 43, 387–418. https://doi.org/10.1093/sysbio/43.3.387

Auffenberg, W. (1980). The herpetofauna of Komodo, with notes on adjacent areas. Bulletin of the Florida State Museum, Biological Sciences, 25, 39–156.

Austin, C. C. (1999). Lizards took express train to Polynesia. Nature, 397, 113–114. https://doi.org/10.1038/16365

Barbour, T. (1912). A contribution to the zoogeography of the East Indian Islands. Memoirs of the Museum of Comparative Zoology, 44, 1–168.

Bouckaert, R., Heled, J., Kühnert, D., Vaughan, T., Wu, C. H., Xie, D., … Drummond, A. J. (2014). BEAST 2: a software platform for Bayesian evolutionary analysis. PLoS Computational Biology, 10(4), e1003537. https://doi.org/10.1371/journal.pcbi.1003537

Boulenger, G. A. (1897). LII.—A list of the reptiles and batrachians collected by Mr. Alfred Everett in Lombok, Flores, Sumba, and Savu, with descriptions of new species. Journal of Natural History, 19, 503–509.

Brown, R. M., Linkem, C. W., Siler, C. D., Sukumaran, J., Esselstyn, J. A., Diesmos, A. C., … Grismer, L. (2010). Phylogeography and historical demography of Polypedates leucomystax in the islands of Indonesia and the Philippines: Evidence for recent human-mediated range expansion?. Molecular Phylogenetics and Evolution, 57, 598–619. https://doi.org/10.1016/j.ympev.2010.06.015

Burbrink, F. T., Lawson, R., & Slowinski, J. B. (2000). Mitochondrial DNA phylogeography of the polytypic North American rat snake (Elaphe obsoleta): a critique of the subspecies concept. Evolution, 54, 2107–2118. https://doi.org/10.1111/j.0014-3820.2000.tb01253.x

Capinha, C., Essl, F., Seebens, H., Moser, D., & Pereira, H. M. (2015). The dispersal of alien species redefines biogeography in the Anthropocene. Science, 348, 1248–1251. https://doi.org/10.1126/science.aaa8913

Darriba, D., Taboada, G. L., Doallo, R., & Posada, D. (2012). jModelTest 2: more models, new heuristics and parallel computing. Nature Methods, 9, 772. https://doi.org/10.1038/nmeth.2109

David, P., Vogel, G., & Vidal, N. (2003). On Trimeresurus fasciatus (Boulenger, 1896) (Serpentes: Crotalidae), with a discussion on its relationships based on morphological and molecular data. Raffles Bulletin of Zoology, 51, 149–158.

Edgar, R. C. (2004). MUSCLE: multiple sequence alignment with high accuracy and high throughput. Nucleic Acids Research, 32, 1792–1797. https://doi.org/10.1093/nar/gkh340

Evans, B. J., Brown, R. M., McGuire, J. A., Supriatna, J., Andayani, N., Diesmos, A., … Cannatella, D. C. (2003). Phylogenetics of fanged frogs: testing biogeographical hypotheses at the interface of the Asian and Australian faunal zones. Systematic Biology, 52, 794–819. https://doi.org/10.1093/sysbio/52.6.794

Ferreira, J.B. (1898). Reptis de Timôr no Museu de Lisboa. Jornal de Sciencias, Mathematicas, Physicas e Naturaes (Segunda Série), 5, 151–156.

Fritts, T. H. (1993). The common wolf snake, Lycodon aulicus capucinus, a recent colonist of Christmas Island in the Indian Ocean. Wildlife Research, 20, 261–265. https://doi.org/10.1071/WR9930261

Gordon, A. L. (2005). The Indonesian Seas. Oceanography, 18, 14-27.

Hall, R. (2009). Southeast Asia’s changing paleogeography. Blumea, 54, 148–161. https://doi.org/10.3767/000651909X475941

Heinsohn, T. (2003). Animal translocation: long-term human influences on the vertebrate zoogeography of Australasia (natural dispersal versus ethnophoresy). Australian Zoologist, 32, 351–376. https://doi.org/10.7882/AZ.2002.014

How, R. A., Schmitt, L. H., & Suyanto, A. (1996). Geographical variation in the morphology of four snake species from the Lesser Sunda Islands, eastern Indonesia. Biological Journal of the Linnean Society, 59, 439–456. https://doi.org/10.1111/j.1095-8312.1996.tb01476.x

How, R. A., & Kitchener, D. J. (1997). Biogeography of Indonesian snakes. Journal of Biogeography, 24, 725–735. https://doi.org/10.1046/j.1365-2699.1997.00150.x

Ingram, G. (1987). Does the skink, Mabuya multifasciata, occur in Australia? Northern Territory Naturalist, 10, 11–12.

Van Kampen, P. N. (1923). The amphibia of the Indo-Australian Archipelago. E. J. Brill, Ltd, Leiden, 304 pp.

Kaiser, H., Carvalho, V. L., Freed, P., & O’Shea, M. (2010). A widely traveled turtle: Mauremys reevesii (Testudines: Geoemydidae) in Timor-Leste. Herpetology Notes, 3, 93–96.

Kraus, F. (2015). Impacts from invasive reptiles and amphibians. Annual Review of Ecology, Evolution, and Systematics, 46, 75–97. https://doi.org/10.1146/annurev-ecolsys-112414-054450

Kuraishi, N., Matsui, M., Hamidy, A., Belabut, D. M., Ahmad, N., Panha, S., … Thong, H. T. (2013). Phylogenetic and taxonomic relationships of the Polypedates leucomystax complex (Amphibia). Zoologica Scripta, 42, 54–70. https://doi.org/10.1111/j.1463-6409.2012.00562.x

de Lang, R. (2011). The snakes of the Lesser Sunda Islands (Nusa Tenggara), Indonesia: a field guide to the terrestrial and semi-aquatic snakes with identification key. Chimaira, Frankfurt am Main, Germany.

Librado, P., & Rozas, J. (2009). DnaSP v5: a software for comprehensive analysis of DNA polymorphism data. Bioinformatics, 25, 1451–1452. https://doi.org/10.1093/bioinformatics/btp187

MacArthur, R. H., & Wilson, E. O. (2001). The theory of island biogeography (Vol. 1).Princeton University Press.

Malhotra, A., & Thorpe, R. S. (2004). A phylogeny of four mitochondrial gene regions suggests a revised taxonomy for Asian pitvipers (Trimeresurus and Ovophis). Molecular Phylogenetics and Evolution, 32, 83–100. https://doi.org/10.1016/j.ympev.2004.02.008

Mertens, R. (1927). Herpetologische Mitteilungen XVII, Mabuya multifasciata, Kuhl auf Bali. Senckenbergiana, 9, 178–181.

Mertens, R. (1930) Die Amphibien and Reptilien der inseln Bali, Lombok, Sumbawa and Flores. Abhandlungen Senckenbergischen Naturforschenden Gesellschaft, 42, 115–344.

Meshaka, W. E., Butterfield, B. P., & Hauge, J. B. (2004). The exotic amphibians and reptiles of Florida. Krieger Pub. Co., Malabar, Fla.

Monk, K. A., De Fretes, Y., & Reksodiharjo-Lilley, G. (1997). The ecology of Nusa Tenggara and Maluku. Singapore: Periplus Editions.

O’Connor, S., Ono, R., & Clarkson, C. (2011). Pelagic fishing at 42,000 years before the present and the maritime skills of modern humans. Science, 334, 1117–1121. https://doi.org/10.1126/science.1207703

Orme, C. D. L., Davies, R. G., Burgess, M., Eigenbrod, F., Pickup, N., Olson, V. A., … Stattersfield, A. J. (2005). Global hotspots of species richness are not congruent with endemism or threat. Nature, 436, 1016. https://doi.org/10.1038/nature03850

O’Shea, M., Kusuma, K. I., & Kaiser, H. (2018). First record of the Island Wolfsnake, Lycodon capucinus, from New Guinea, with comments on its widespread distribution and confused taxonomy, and a new record for the Common Sun Skink, Eutropis multifasciata. IRCF Reptiles and Amphibians, 25, 70–84. http://hdl.handle.net/2436/621615

Reilly, S. B. (2016). Historical biogeography of reptiles and amphibians from the Lesser Sunda Islands of Indonesia. Doctoral Dissertation, University of California, Berkeley, USA.

Reilly, S. B., Wogan, G. O., Stubbs, A. L., Arida, E., Iskandar, D. T., & McGuire, J. A. (2017). Toxic toad invasion of Wallacea: A biodiversity hotspot characterized by extraordinary endemism. Global Change Biology, 23, 5029–5031. https://doi.org/10.1111/gcb.13877

Reilly, S. B., Stubbs, A. L., Karin, B. R., Bi, K., Arida, E., Iskandar, D. T., & McGuire, J. A. (2019). Leap-frog dispersal and mitochondrial introgression: phylogenomics and biogeography of Limnonectes fanged frogs in the Lesser Sundas archipelago of Wallacea. Journal of Biogeography, https://doi.org/10.1111/jbi.13526

Roesler, H., Bauer, A. M., Heinicke, M. P., Greenbaum, E., Jackman, T., Nguyen, T. Q., & Ziegler, T. (2011). Phylogeny, taxonomy, and zoogeography of the genus Gekko Laurenti, 1768 with the revalidation of G. reevesii Gray, 1831 (Sauria: Gekkonidae). Zootaxa, 2989, 1– 50.

De Rooij, N. (1917a). The Reptiles of the Indo-Australian Archipelago: Lacertilia, Chelonia, Emydosauria (Vol. 1). EJ Brill.

De Rooij, N. (1917b). The Reptiles of the Indo-Australian Archipelago: Ophidia (Vol. 2). EJ Brill.

Schmitt, L. H., How, R. A., Hisheh, S., Goldberg, J., & Maryanto, I. (2000). Geographic patterns in genetic and morphological variation in two skink species along the Banda Arcs, southeastern Indonesia. Journal of Herpetology, 34, 240–258. doi:10.2307.1565421

Smith, M. J., Cogger, H., Tiernan, B., Maple, D., Boland, C., Napier, F., … & Smith, P. (2012). An oceanic island reptile community under threat: the decline of reptiles on Christmas Island, Indian Ocean. Herpetological Conservation and Biology, 7, 206–218.

Stamatakis, A. (2014). RAxML version 8: a tool for phylogenetic analysis and post-analysis of large phylogenies. Bioinformatics, 30, 1312–1313. https://doi.org/10.1093/bioinformatics/btu033

Tajima, F. (1989). Statistical method for testing the neutral mutation hypothesis by DNA polymorphism. Genetics, 123, 585–595.

van Lidth de Jeude, T. W. (1895). Reptiles from Timor and the neighbouring islands. Notes from the Leyden Museum, 16, 119–127.

Whittaker, R. J., & Fernández-Palacios, J. M. (2007). Island biogeography: ecology, evolution, and conservation. Oxford University Press.

Whitten, A. J., Mustafa, M., & Henderson, G. S. (1987). The Ecology of Sulawesi. Gadjah Mada University Press, Yogyakarta.

Yuwono, F. B. (1998). The trade of live reptiles in Indonesia. Mertensiella, 9, 9-15.

